# Genomic contacts reveal the control of sister chromosome decatenation in *E. coli*

**DOI:** 10.1101/2021.05.17.444411

**Authors:** Brenna Conin, Ingrid Billault-Chaumartin, Hafez El Sayyed, Charlotte Cockram, Romain Koszul, Olivier Espéli

## Abstract

In bacteria, chromosome segregation occurs progressively, from the origin to the terminus, a few minutes after the replication of each locus. In-between replication and segregation, sister loci are maintained in an apparent cohesive state by topological links. Whereas topoisomerase IV (Topo IV), the main bacteria decatenase, controls segregation, little is known regarding the influence of the cohesion step on chromosome folding. In this work, we investigated chromosome folding in cells with altered decatenation activities. Within minutes after Topo IV inactivation, a massive chromosome reorganization takes place, associated with increases in trans-contacts between catenated sister chromatids and in long-range cis-contacts between the terminus and distant loci on the genome. A genetic analysis of these signals allowed us to decipher specific roles for Topo IV and Topo III, an accessory decatenase. Moreover we revealed the role of MatP, the terminus macrodomain organizing system and MukB, the *E. coli* SMC in organizing sister chromatids tied by persistent catenation links. We propose that large-scale conformation changes observed in these conditions reveal a defective decatenation hub located in the terminus area. Altogether, our findings support a model of spatial and temporal partition of the tasks required for sister chromosome segregation.

## Introduction

Prokaryotic and eukaryotic chromosomes are not randomly folded, but consist of well-defined structural entities with a complex hierarchical organization. The regulation of this network and its functional interplay with gene expression or other chromosomal metabolic processes such as DNA repair, replication and segregation have been actively investigated in a number of species (Wang and Rudner, 2014). Improvement in imaging techniques of living cells, as well as the development of genomic approaches such as chromosome conformation capture (3C/Hi-C; reviewed in Dekker et al., 2013) techniques have highlighted a mosaic of intertwined structural features including loops, domains, and compartments (Dame et al., 2020).

In bacteria, an important part of genome folding relies on the presence of free DNA supercoils. Independent supercoiling domains were first observed in the 1970’s through a combination of molecular biology and electron microscopy approaches (Kavenoff and Bowen, 1976; Sinden and Pettijohn, 1981). These textbook pictures consist of a succession of large (~200 kb) DNA plectonemes, or microdomains, that are delimited by topological insulators. Genomic and recombination assays confirmed the existence of these microdomains, but found that they are likely smaller in size (10-50 kb), delimited by stochastic barriers (Postow, 2004; Stein et al., 2005). These results suggest that microdomains are not static, and that their size and position can be modulated by DNA transactions such as transcription (Deng et al., 2005) or the binding of proteins to DNA (Leng et al., 2011). The democratization of 3C/Hi-C methodologies provided the opportunity to further assess the impact of DNA supercoiling on chromosome folding. To date, 3C/Hi-C data revealed the organization of large chromosomal features, such as the alignment of replication arms (Le et al., 2013; Marbouty et al., 2015; Wang et al., 2015), macrodomains (Lioy et al., 2018) or inter-chromosomal contacts (Val et al., 2016). At smaller scale, Hi-C contact maps of every bacterial genome studied to date display self-interacting domains (CIDs, 20-200kb), whose significance remain unclear (Böhm et al., 2020; Le et al., 2013; Lioy et al., 2018; Marbouty et al., 2015; Val et al., 2016; Wang et al., 2015). The boundaries in-between CIDs correlate with the presence of long, highly expressed genes, or genes coding membrane proteins (Le and Laub, 2016; Le et al., 2013; Lioy et al., 2018).

Simulations suggest that supercoils are able to organize bacterial genome because they would condense DNA and promote the disentanglement of topological domains (Holmes and Cozzarelli, 2000). DNA supercoiling is under tight homeostatic control by topoisomerases. In *E. coli*, Topoisomerase I (Topo I), Topoisomerase III (Topo III), DNA Gyrase and Topoisomerase IV (Topo IV) have all well characterized enzymatic activities (Champoux, 2001; Levine et al., 1998), but understanding the roles played by topoisomerases in chromosomal compaction, folding and organization is still a work in progress. DNA Gyrase, that promotes the formation of free supercoils, is the best candidate for regulation of chromosome organization through supercoiling. Hi-C analysis following gyrase inhibition was studied in *Caulobacter crescentus*, revealing modest changes in chromosome conformation with a slight decrease of 20- to 200-kb contacts that reduced the sharpness and positions of CID boundaries (Le et al., 2013). This observation agrees with recombination data showing that topoisomerase alterations reduce supercoiling domain’s size (Staczek and Higgins, 1998). The contribution of other topoisomerases in establishing, maintaining, and regulating genome-wide DNA contacts has, so far, not been investigated.

The *E. coli* SMC complex MukBEF was shown to be linked to DNA supercoiling, with MukBEF defects being suppressed by Topo I mutation (Sawitzke and Austin, 2000). The suppression correlates with an excess of negative supercoiling promoted by DNA gyrase. In addition, Hi-C analysis of a MukB mutant showed an important loss of long-range contacts (Lioy et al., 2018a), suggesting that MukB promotes contacts between distant pairs of loci. One hypothesis is that MukB could promote DNA loops *in vivo* (Badrinarayanan et al., 2012; Baxter et al., 2019; Carter and Sjögren, 2012; Ruiten and Rowland, 2018). For instance, it is possible that MukB, organised in an axial core, extrudes plectonemic microdomains (Mäkelä and Sherratt, 2020). Interestingly, MukB interacts with ParC, the catalytic subunit of Topo IV. Their interaction changes the catalytic properties of Topo IV (Hayama and Marians, 2010; Li et al., 2010) and modulates its localization (Nicolas et al., 2014; Stracy et al., 2015). *In vitro* assays suggest that a MukB – Topo IV interaction promotes DNA compaction by forcing the intramolecular knotting activity of Topo IV (Kumar et al., 2017). Single molecule imaging reveals that a small portion of ParC molecules are associated with MukB in small clusters, (Stracy et al., 2015). The absence of MukB leads to a 2-fold reduction of Topo IV clusters, suggesting that MukB is a driver for the localization of Topo IV in the cell

Topo IV is the main bacterial decatenase. It catalyzes the elimination of precatenanes and catenanes formed during replication (Zechiedrich et al., 1997), and is essential for proper chromosome segregation (Joshi et al., 2013; Kato et al., 1990; Lesterlin et al., 2012; Wang et al., 2008) (Kato 1990). The requirement of PriA, the main replication fork restart protein, for the survival of Topo IV thermosensitive mutants (*parE^ts^* and *parC^ts^*), suggests that Topo IV inactivation increases replication fork stalling (Grompone et al., 2004). Nevertheless, chromosome replication in *parE^ts^* mutant cells occurs at a similar rate to that seen in *WT* cells (Wang et al., 2008). Topo IV activity is highest at the *dif* site positioned in the middle of the Terminus of replication macrodomain, presumably to solve catenanes (El Sayyed et al., 2016; Hojgaard et al., 1999; Mercier et al., 2008). However, it also presents hundreds of putative activity sites dispersed on the genome, presumably used for the removal of precatenanes (El Sayyed et al., 2016).

Here we investigate, using Hi-C, the effect of the different topoisomerases on the *E. coli* chromosome conformation. The inactivation of Topo IV generates the most significant changes in the organization of the chromosome, with a pair of long-range contact patterns, hereafter called butterfly wings, expanding from the *terminus* macrodomain flanking regions. Imaging experiments confirmed the chromosome reorganization when Topo IV activity is reduced, suggesting that in these conditions distant sister loci segregate when they contact the terminus. Because MatP and MukB also influence the butterfly pattern, we propose that it reveals a particular genome folding dedicated to decatenation. We also showed that when Topo IV is deficient, Hi-C reveals inter-molecular contacts between sister chromosomes. Topo III limits these contacts suggesting that it eliminates an important part of precatenanes. Together, our results highlight the ability of Hi-C to unveil chromosomal topological features including inter-chromosomal contacts, to improve our understanding of the mechanisms governing chromosome segregation.

## Results

### Topo IV inactivation induces large-scale chromosome conformation changes in enterobacteria

To investigate the respective contribution of each topoisomerase to the genome-wide organization of bacteria chromosomes, we applied capture of chromosome conformation (Hi-C) (Cockram et al., 2021; Lieberman-Aiden et al., 2009); Material and Methods) to exponentially growing *Escherichia coli* cells either mutated or inactivated for each of the four topoisomerases. The ratio between *wt* and mutant/depleted cells contact maps were plotted (Material and Methods) (Lioy et al., 2018).

The inactivation of Topo IV using thermosensitive alleles of either the *parC* (*parC^ts^*) or *parE* (*parE^ts^*) subunits (Kato et al., 1990) resulted in significant changes in the global chromosomal architecture (**Figure 1A and 1B**). At the permissive temperature (30°C), the *parE*^ts^ mutant grew similarly to wt, with no significant defects in cell growth nor changes in chromosome conformation (**Supplementary Figure 1A** and **1C**, respectively). Following a 1-hour shift to non-permissive temperature (42°C), we observed changes in mid- to long-range DNA contacts at three regions of the genome: First, an increase in short- to medium-range contacts from *oriC* to positions located at ~1 Mb and ~2 Mb on the right and left replichores, respectively. Second, a strong reduction of mid-range contacts in the terminus region (Coordinates 1,315 kb −1,830 Kb, **Figure 1A-B**) accompanied by an increase of very short-range (< 50kb, 10 bins) contacts. This particular terminus pattern also featured a strong CID-like border around the *dif* site. Finally, a peculiar, butterfly-like signal, with two wings of long-range contacts emerges from the flanking regions of the Ter, and extending towards *oriC*.

**Figure 1:**
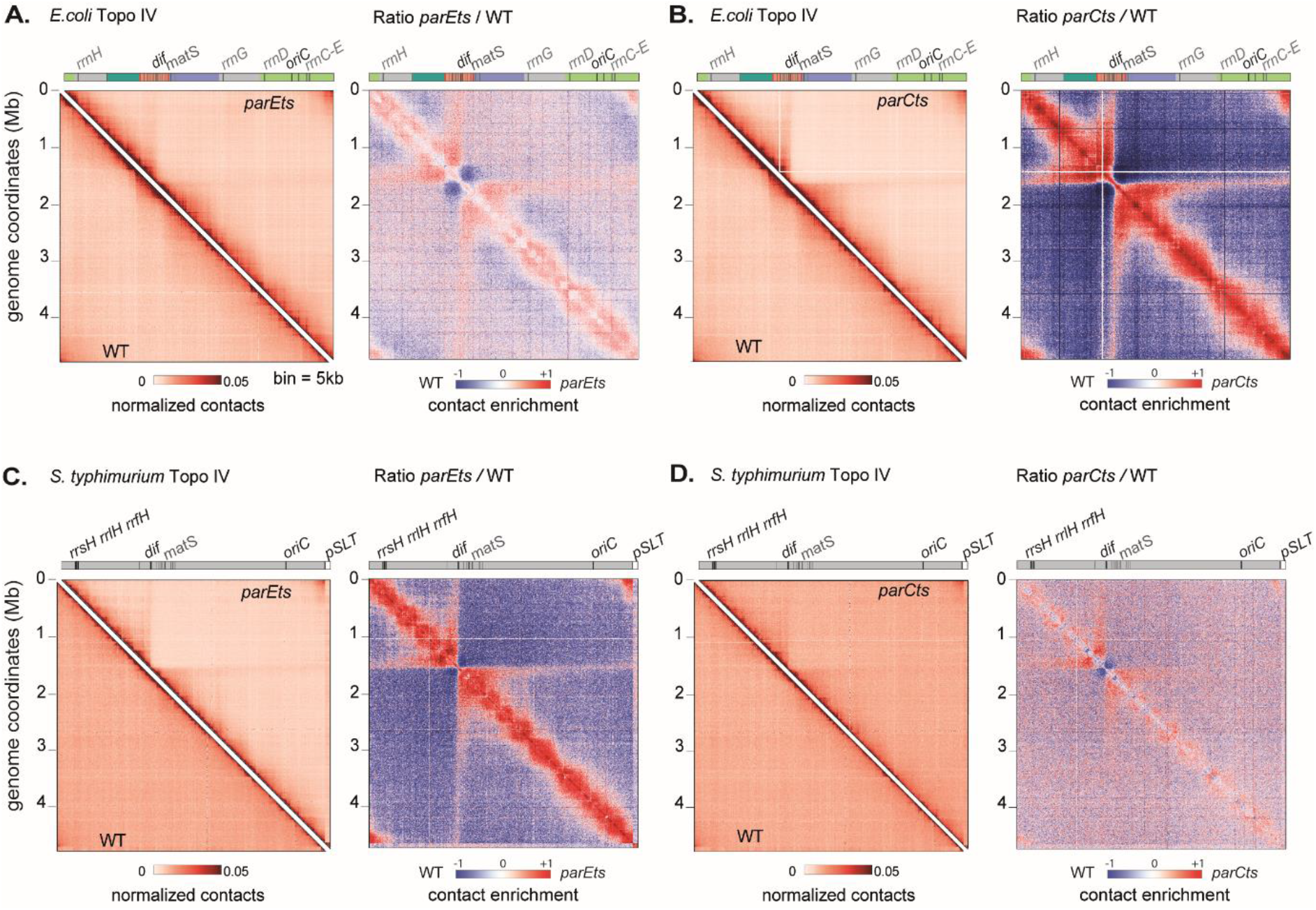
Impact of Topoisomerase IV alteration on nucleoid organization of E. coli and S. typhimurium. For each panels, symmetric halves of the normalized contact map binned at 5kb with the wild type (WT) on the bottom and the altered topoisomerase on the top, and the corresponding ratio matrix. (A) parE^ts^ after 60min of shift to non-permissive temperature (42°C) in E. coli, (B) parC^ts^ grown at 30°C in E. coli, (C) parE^ts^ after 60min of shift to non-permissive temperature (42°C) in S. typhymurium, (D) parC^ts^ after 60min of shift to non-permissive temperature (42°C) in S. typhymurium. Genome coordinates are indicated by the x and y axes. Interesting positions of the genome are indicated above the plot. matS sites are represented as grey bars. For E. coli, macrodomains are represented by light green (ori), dark green (right), red (ter), blue (left), gray (NR/NL). For the normalized contact maps, the color scale of the frequency of contacts between two regions of the genome is indicated below (arbitrary units), from white (rare contacts) to dark red (frequent contacts). For the ratio matrices, a decrease or increase in contacts in the mutant cells compared with the control is represented with a blue or red color, respectively. White indicates no differences between the two conditions.

These changes in chromosome folding are rescued by a plasmid expressing ParE into the *parE*^ts^ mutant cells **(Supplementary Figure 1B-1D)**. Although *parC^ts^* mutants are sicker, even at low temperatures **(Supplementary Figure 1A)**, we observed a phenotype similar to that of *parE^ts^*, suggesting that the chromosomal changes observed in these mutants are associated with Topo IV inactivation **(Figure 1B)**. To broaden these observations, we applied Hi-C to another y-proteobacteria, *Salmonella typhimurium* for which thermosensitive Topo IV mutations are available (Luttinger et al., 1991; Springer and Schimd, 1993). Similar conformation changes were observed **(Figure 1C and 1D).**

Depletion, deletion or inactivation of the three others *E. coli* topoisomerases (Topo I, Topo III and gyrase) had only mild effect on the global *E. coli* chromosome organization (**Supplementary Figure 1E, 1F and 1G).** To reduce Topo I activity we used the topA31 allele (Conter et al., 1997), a *topB* deletion mutant was used to study Topo III (DiGate and Marians, 1989), and DNA gyrase was studied using a thermosensitive mutant of *gyrB* (Orr and Staudenbauer, 1981). We observed that although alterations in Topo I and Topo III resulted in a slight loss or gain in short-range contacts, no other significant changes in the chromosome conformation of these cells were detected **(Supplementary Figure 1E and 1F)**.

Since gyrase inhibition blocks DNA replication and results in a significant remodeling of the transcriptome (Orr and Staudenbauer, 1981; Peter et al., 2004), *gyrB^ts^* mutant cells were only shifted to the non-permissive temperature for a brief (20 min) period. Cells exhibit an altered chromosomal organization compared to *WT* cells growing at the same temperature **(Supplementary Figure 1G)**. Inhibition of Gyrase activity resulted in a decrease of short-range contacts. These observations agree with the involvement of gyrase in the definition of supercoiling microdomains or CIDs (Le et al., 2013).

### Hi-C features observed in *parE*^ts^ mutant cells are concomitant with a large reorganization of the nucleoid

Imaging further supported that nucleoid organization and compaction undergoes a progressive change following *parE^ts^* inactivation. Following the 42°C shift, the average DAPI staining density roughly doubled compared to *wt* **(Figure 2A-B)**. Using *parS* pMT1 tags in the *oriC* and *terminus* region, we analyzed the segregation of sister foci after replication **(Figure 2C).** In agreement with earlier publications (Wang et al., 2015),we observed a reduction in foci number after 40 min at non-permissive temperature, suggesting that the increase in nucleoid compaction correlates with the blockage of sister chromatid segregation. In addition, we observed a strong change in the localization of *oriC* and *ter* foci within the cell (**Figure 2D and 2E and Supplementary Figure 2A and 2B**). At permissive temperature, *oriC* positions preferentially towards the nucleoid edges whereas the *ter* positions closer to the center of the nucleoid. Upon shift to non-permissive temperature, this organization is flipped, with the *ter* and *oriC* migrating towards the cell pole and mid-cell, respectively. These observations suggest that Topo IV inactivation affects both sister chromosome segregation and nucleoid organization.

**Figure 2:**
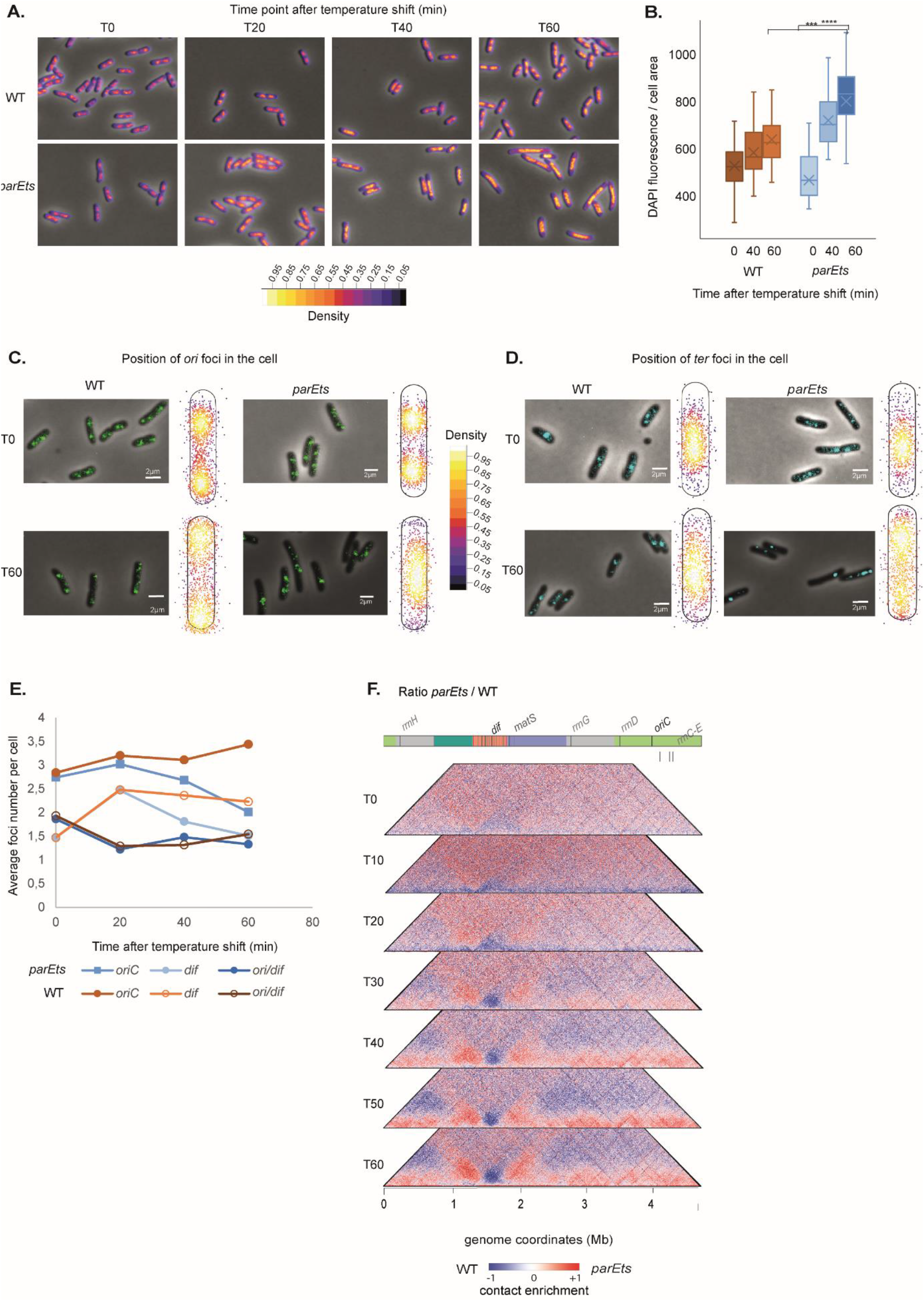
Hi-C features observed in *parE^ts^* mutant cells are concomitant with a large reorganization of the nucleoid. (A) Merged images of phase contrast and DAPI signal (DIC signal in grey and the density of the DNA signal is represented with a fire color scale) of WT (left) and parE^ts^ (right) cells every 20min during a temperature shift. Scale bar = 2 μm. (B) Quantification of the fluorescence density as proxy for DNA density across time of the WT (brown boxes) and parE^ts^ (blue boxes) (a Student test, *** P <0.0005). (C-D) Merged images of phase contrast and GFP signal, and average position of the foci in WT (left) and parE^ts^ (right) cells before temperature shift (T0) and after 60min shift (T60). In (C), GFP signal reveals the position of aidB parS T1 locus (ori) in the cells. In (D), GFP signal reveals the position of fear parS T1 (ter) in the cells. Dots represents each detected focus (n=200) and the color scale represents the density of detected foci in the cell from black (0.05) to white (1). Intermediate time analysis can be found in Supplementary Figure 2A and 2B. (E) Quantification of the number of ori and ter foci across the time course as a proxy for sister chromatids cohesion, and of the ratio ori/ter for WT (brown lines) and parE^ts^ (blue lines (F) Kinetics of the impact of Topo IV alteration with a time point every 10min after shift at non permissive temperature from t0 to t60 represented by the ratio matrices of parE^ts^ cells versus WT cells kinetics. Normalized contact map of each time point and t120 can be found in Supplementary Figure 2C-F. matS sites are represented as gray bars. Macrodomains are represented by light green (ori), dark green (right), red (ter), blue (left), gray (NR/NL). The y axis indicates the genomic coordinates (Mb). A decrease or increase in contacts in the mutant cells compared with the control is represented with a blue or red colour, respectively. White indicates no differences between the two conditions.

We then performed a kinetic analysis, using Hi-C, of chromosome folding after Topo IV inactivation. It allowed us to evaluate whether nucleoid conformation changes correlate with the apparition of new contacts in the Hi-C maps **(Figure 2F and Supplementary Figure 2C)**. The first consequence of Topo IV inactivation was an immediate loss of short-range contacts over the entire genome **(Figure 2F, T10).** This drop was gradually substituted by an increase in short- to mid-range contacts, which becomes prevalent along chromosome arms after 40 minutes. Concomitantly the butterfly wings expanded from positions 1Mb and 2Mb. These features correlated with the evolution of the nucleoid density and organization over the same time period. The *terminus* region behaved differently, showing a persistent lack of short-range interactions over the entire kinetics. We could not determine whether this terminus pattern was a cause or a consequence of its localization at the nucleoid periphery (**Figure 2E)**. After two hours (t120), we observed similar features with an overall increase of the differences with the *wt* **(Supplementary Figure 2D-G).** Altogether, these time course experiments show that specific contacts accumulate at discrete positions surrounding the *ter* domain upon inactivation of Topo IV, and therefore that Topo IV plays a role in suppressing the formation of long-range contacts within or between sister chromosomes.

### The absence of Topo IV activity results in the co-localization of distant loci with the terminus of the chromosome

We characterized further the two butterfly-wing stripes emanating at a 45° angle from the edges of the *terminus* region in the ratio contact maps between Topo IV and *wt* **(Figure 3A** coordinates 1.3 − 1.5 Mb and 1.5 − 1.8Mb, respectively). These signals correspond to an enrichment in medium and long-range contacts by ~200 kb regions that in one direction span all the way up to *oriC*, while on the other direction, are blocked by *dif*, which represents a barrier preventing contacts across each halves of the *terminus* **(Figure 3A and B)**. Using a 4C-like plot, we investigated how specific 5kb bins contribute to the butterfly signal (**Figure 3C and D**). As illustrated for two representative regions (position 3,010 and 4,440 Kb), a small (≈ 1.5 fold), but significant, contact enrichment with the *dif* zone was detected several megabases away from *dif*. This enrichment involved several (3 to 10) bins positioned a few tens of kb away from *dif.* The contacts between the *dif* zone and the 3,010 kb or 4,440 kb regions present a comparable frequency, suggesting that they do not obey to genomic distance law. We observed, on zoomed panels, that the bin containing *dif* (green dot on Figure 3D) was not involved in the butterfly wing contacts **(Figure 3D).** To test this exclusion of *dif,* we measured the sum of contacts made by a 100 kb window 2Mb away from *dif* (Σ sign of **Figure 3A**) with the 1.2Mb region containing *dif* **(Figure 3E).** This window displayed enriched contacts with most bins from the *terminus* region over 200 kb surrounding *dif*, but not with *dif* itself. Their distance independent frequency and the lack of asymmetry at *dif* strongly suggests that butterfly wings correspond to long-range 3D contacts rather than slithering events.

**Figure 3:**
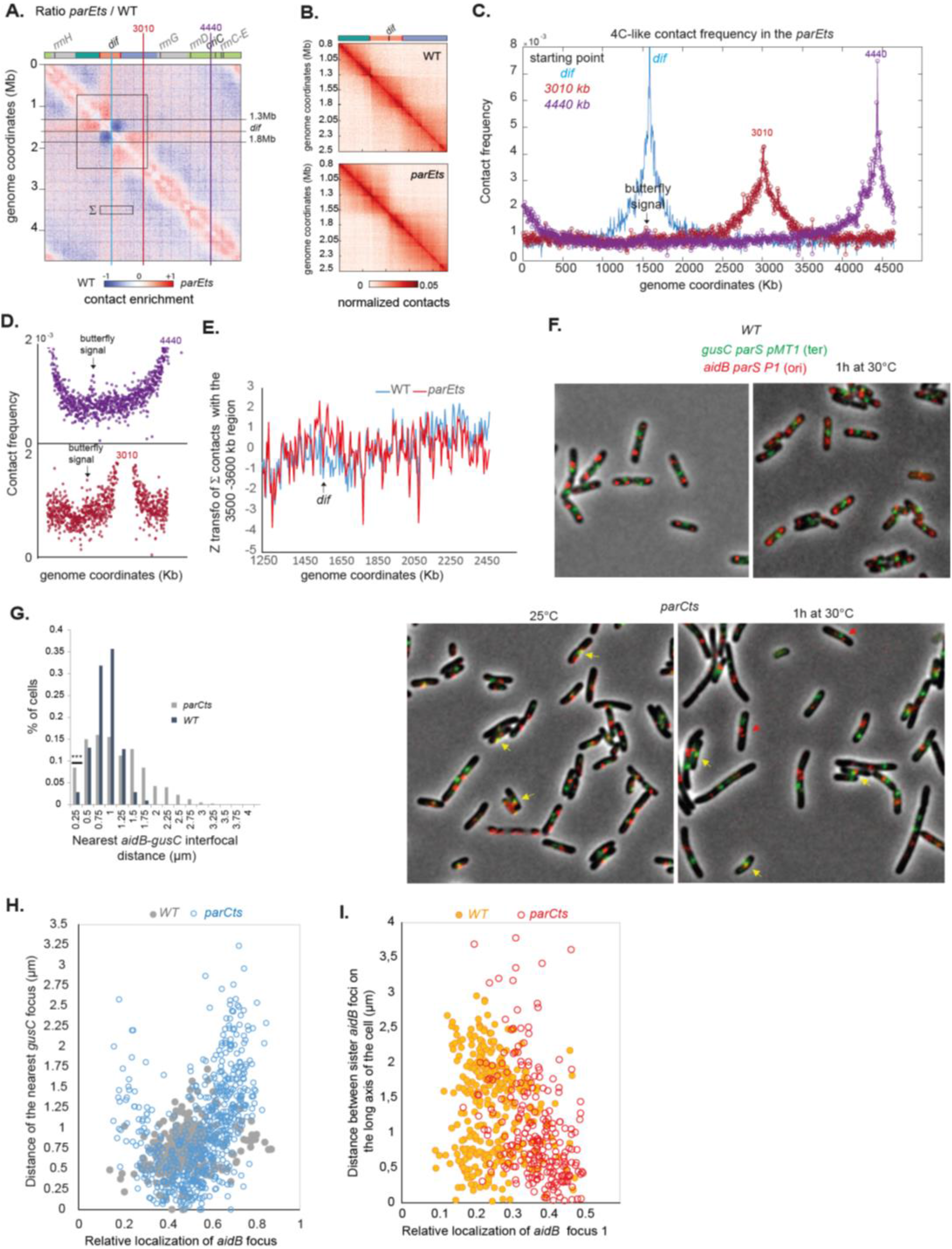
Characterization of the butterfly wings pattern. (A) Ratio of the normalized contact map of parE^ts^ and WT where the butterfly wings borders are indicated by gray lines (1.3Mb – dif and dif − 1.8Mb) and two representative zones at 3301kb (red) and 4440kb (purple). (B) Zoom on the terminus macrodomain of the WT and parE^ts^ matrices from 0.8Mb to 2.5Mb (C) 4C-like plot of contacts from the dif, the 3010Kb and 4440 Kb positions, (D) Rescaling of the plot C to show the butterfly wing contacts in the dif area. (E) Z transformation of the contacts made by a 100 Kb region located between the coordinates 3500 Kb and 3600 Kb (marked Σ in A) with the terminal part of the genome (coordinates 1250 - 2450 Kb). WT and parE^ts^ normalized matrix were analyzed. (F) Merged images of phase contrast, CFP and Y-GFP signal. Y-GFP signal reveals the position of gusC parS pMT1 (ter) in the cells and CFP signal reveals the position of aidB parS P1 locus (ori) in the WT (left) and parC^ts^ cells (right) at permissive temperature (25°C) and after 1h at non permissive temperature (30°C). Yellow arrows indicate colocalisation between aidB and gusC foci and red arrows indicate close sister aidB foci at the center of the cells. Scale bars are 4 μm. (G) Distribution of the nearest interfocal distance between ter and ori foci in the WT and parC^ts^ strain after 1h at 30°C. (N= 600; colocalization = IFD <250nm) was analyzed with a KS test, *** P <10^−9^). (H) Analysis of nearest interfocal distance between ter and ori foci as a function of the relative localization of the ori focus after 1h at 30°C. (I) Analysis of the distance between sister ori foci as a function of the position of the closest ori focus to the pole after 1h at 30°C.

To confirmed the propagation of these very long-range contacts at the single cell level, the inter-focal distances between fluorescently labelled loci positioned at *ori* and were measured in cells carrying the *parC^ts^* allele (**Figure 3F-I**), that displayed butterfly wings at low temperature (30°C) (**Figure 1C**). In these conditions, the *ori* (*aidB,* 4.4Mb) and the *ter* (*gusC*, 1.69Mb) loci colocalized at the center of the cell in approximately 8% of cells, a proportion significantly higher than for the *wt* **(Figure 3G)**. The accumulation of the *ori* region at the center of the cell agrees with the global nucleoid rearrangement observed upon Topo IV inactivation **(Figure 2)**. Ori-ter colocalization was observed close to mid-cell, while most *ter* foci were localized near the pole of the nucleoid **(Figures 2D and 3H)**. Interestingly, the closest distances between sister *ori* foci were also observed close to mid-cell in the *parC^ts^* strain **(Figure 3I).** This was in sharp contrast with *wt* where the minimal distance between sister *ori* foci was typically observed at the ¼ and ¾ positions. This observation suggests that when Topo IV is inactivated, the release of *ori* pairing takes place closer to mid-cell. Imaging therefore confirms the existence of occasional long-range contacts between the *terminus* and distant regions of the chromosome in absence of Topo IV activity, and suggests that these contacts may correspond to sister loci segregation attempts. The detection of rare *ori-ter* colocalization events in *wt* cells **(Figure 3G)** suggests that the peculiar chromosome folding events detected in the absence of Topo IV activity might also exist in some *wt* cells.

### *parE^ts^* inactivation increase inter-chromosomal contacts

The ability of conventional Hi-C to capture intra-chromosomal (*cis*) contacts is well established. However, its ability to reveal *trans* contacts between sister-chromatids remains limited given the difficulty to distinguish both homologous molecules without incorporation of modified bases (Oomen et al., 2020). Since Topo IV is mainly known for its involvement in the removal of inter-chromosomal links (or catenanes), presumably between allelic (or near-allelic) loci, we sought to test whether the emergence of prominent Hi-C signals spanning throughout the *parE*^ts^ contact map could result from *trans* contacts. We therefore sought to distinguish the contribution of intra vs. inter-molecular contacts to these patterns by determining the impact of DNA replication in absence of Topo IV activity. To do this, we blocked replication initiation by treating WT and *parE^ts^* cells with the amino acid analog dl-serine hydroxamate (Ferullo et al., 2009) (**Figure 4A)**. Following treatment, the cells were then shifted to 42°C for 1 hour. Hi-C ratio maps of replicating vs. non-replicating *wt* cells revealed a significant decrease in mid-range contacts along the entire chromosome in the absence of replication **(Figure 4B)**. Non-replicating *parE*^ts^ cells did not show the characteristic pattern observed for replicating cells **(Figure 4C and Supplementary Figure 3A-B).** Since the stringent response mediated by SHX also reduces transcription of many genes (Sharma and Chatterji, 2010), we tested whether absence of the characteristic pattern in SHX-treated cells could result from a poor transcription. Asynchronously growing *parE*^ts^ and WT cells were therefore treated with the antibiotic rifampicin to inhibit transcription (STAR Methods). Although rifampicin alters Hi-C matrices’ quality (Le et al., 2013), the partition of the chromosome into 3 regions in *parE*^ts^ cells remained visible upon inactivation of Topo IV at 42°C (i.e. the butterfly wings, a higher mid-range contacts in the *oriC* proximal region, and lower mid-range contacts in the *ter*) **(Supplementary Figure 3C and 3D)**. Taken together, these results suggest that Topo IV inactivation only induces characteristic Hi-C patterns in replicating cells, presumably because they correspond to inter-chromosome contacts.

**Figure 4:**
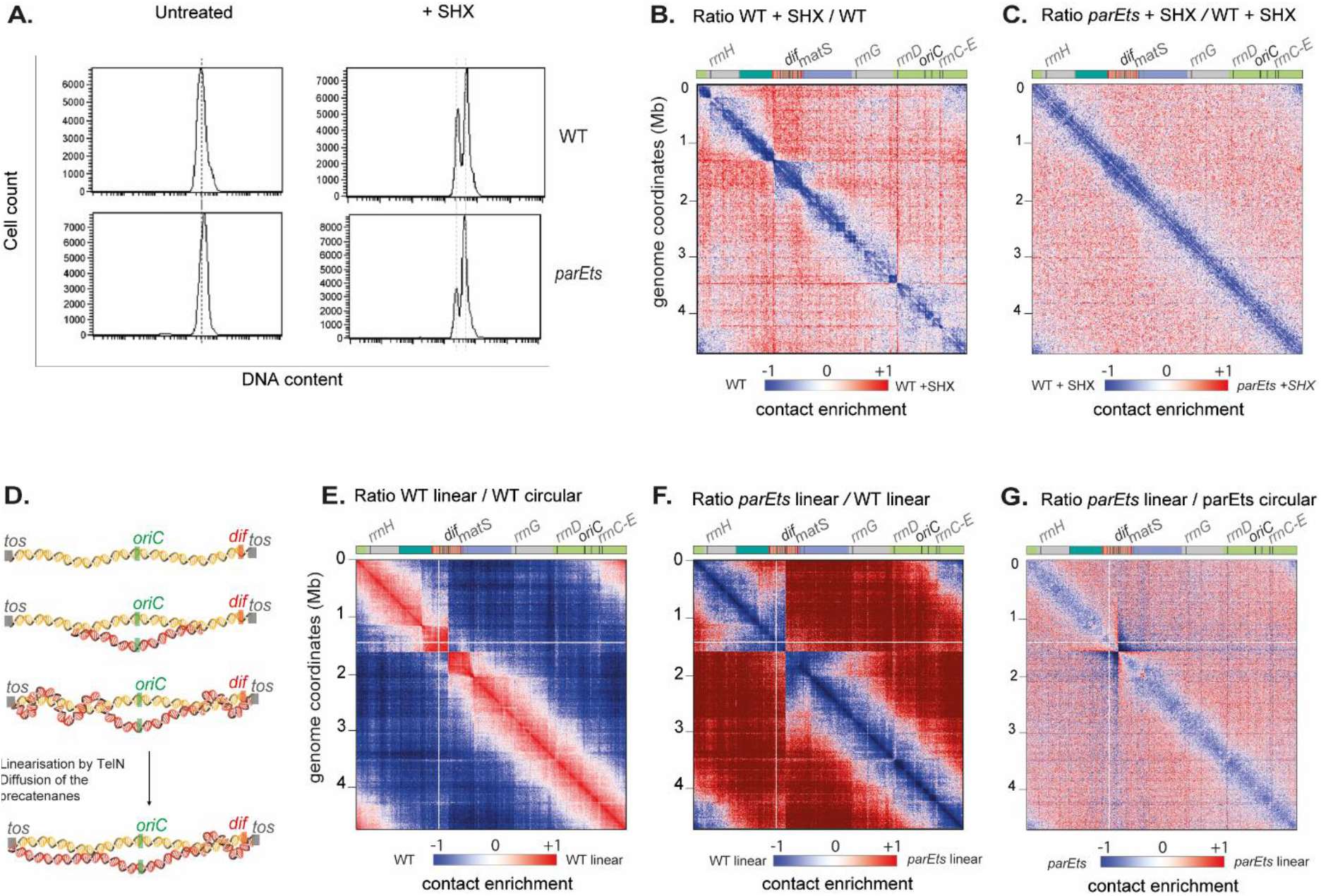
Hi-C features resulting from parE^ts^ alteration result from sister chromosome interactions. (A) FACS of WT and parE^ts^ cells after 60 min of shift at 42°C untreated (left) and treated with 10mg/ml of SHX for 90min total (right). Nucleoid were stained with propidium iodide. Ratio of normalized contacts map binned at 5kb of (B) WT + SHX compared to WT, (C) parE^ts^ + SHX compared to WT + SHX. (D) Graphic representation of the replication in E. coli linear strain. The chromosome is linearized via the tos / TelN system of the phage N15. tos sites (gray box) were inserted in the terminus macrodomain. The replication starts at the oriC site (green box) and progress in a bidirectional manner toward the dif site (red box). We hypothesis that precatenanes are formed during the replication of the linear chromosome in the same manner as in the circular one. Sister chromatids are represented by double helices in red (sister A) and yellow (sister B). Before chromosome segregation, the protelomerase TelN linearize the sister chromosomes at the tos site which could possibly allow for diffusion and resolution of the majority of the precatenanes. Ratio of normalized contacts map binned at 5kb of (E) linear WT vs circular WT, (F) linear parE^ts^ vs linear WT, and (G) linear parE^ts^ linear vs parE^ts^ circular. Macrodomains and interesting positions of the genome are indicated above the plot akin to Figure 1. The y axis indicates the genomic coordinates. A decrease or increase in contacts in the mutant cells compared with the control is represented with a blue or red colour, respectively. White indicates no differences between the two conditions.

### Butterfly wings signals are linked to chromosome topology

We then wondered whether the circular nature of the *E. coli* chromosome could lead to topological constraints on the *terminus* region that would translate into specific contact patterns in the absence of Topo IV. To test this, we performed Hi-C experiments in strains carrying a genome linearized at a *tos* site, inserted near *dif,* thanks to the bacteriophage N15 telomerase (Cui et al., 2007). In this strain, Topo IV is still required for growth (Cui et al., 2007). However, it is no longer active at *dif*, and the filamentation phenotype of the *matP* mutant is now rescued (El Sayyed et al., 2016). These observations hinted that topological tension resulting from Topo IV inactivation freely diffuses when the N15 telomerase linearizes the duplicated chromosome **(Figure 4D)**. In the wt strain, linearization of the chromosome near *dif* resulted in an important redistribution of long-range contacts towards short and mid-range contacts **(Figure 4E and Supplementary Figure 3E)**. Upon inactivation of *parE^ts^*, the Hi-C contact maps of the linear chromosome displayed a significant loss of short- to mid-range contacts, no butterfly wings and no *terminus* pattern compared to circular *wt* chromosomes **(Figure 4F and Supplementary Figure 3F)**. The comparison between *parE^ts^* strains carrying either a linear or a circular genome further revealed a decrease in short/medium contacts along linear chromosomes, in sharp contrast to *wt* conditions, and compatible with a reduction in the number or density of precatenation links **(Figure 4G).** This suggests that the linearization of the strain suppresses the accumulation of interminglements between the sister chromatids. Altogether, these observations strongly support the hypothesis that the enhanced mid-range contacts along chromosome arms, as well as the butterfly wings signals, form in response to a failure of Topo IV to remove inter-chromosomal links.

### Topo III activity partially rescues Topo IV deficiencies on chromosome conformation

Topo III (*topB*) is a type I topoisomerase whose role during the normal cell cycle remains unclear. Both the modest chromosome conformation changes **(Supplementary Figure 1F)** and normal viability observed in the mutant **(Figure 5A)** confirmed that in laboratory growth conditions its role is limited. However, part of its activity is revealed when Topo IV is impaired, as Topo III overexpression rescues *parE^ts^* growth defects (Lee et al., 2019; Nurse et al., 2003**; Figure 5A)**, whereas *topB* deletion reduces the viability of *parE^ts^* strains **(Figure 5A**). When *parE^ts^* was inactivated in a *topB* deletion strain, the Hi-C pattern observed in *parE*^ts^ mutant cells at the non-permissive temperature was strengthened **(Figure 5B)**, with mid-range contacts increase, and stronger butterfly and *terminus* patterns **(Figure 5B).** In the *topB parE^ts^* strain, long-range contacts were observed between distant regions and a large central region of the ter macrodomain **(Figure 5C).** Their frequency was higher **(Figure 5C)** compared to the single *parE^ts^* strain **(Figure 3C).** As observed for the *parE^ts^* strain the *dif* bin appeared to make less long-range contacts than its neighbors **(Figure 5C)**. Interestingly, *parE^ts^* features were visible in the contact maps at 30°C in the absence of Topo III **(Supplementary Figure 4A)**, suggesting that at this temperature Topo IV is already partially inactivated, but that the organization defects are suppressed by Topo III. This is in good agreement with the CFU data **(Figure 5A).** Ratio plot and 4C-like analysis also revealed important changes in the distribution of short and mid-range contacts outside of the *terminus* region when both Topo IV and Topo III are inactivated. When plotting the ratio map between the *parE^ts^ topB* matrix and *topB* or *parE^ts^* matrices we characterized the contribution of Topo III and Topo IV. Butterfly wings, and mid-range contacts appeared in the *parE^ts^ topB* compared to *topB* matrices **(Supplementary Figure 4B-D)**. On the other hand, comparing the *parE^ts^ topB* matrix with *parE^ts^* highlighted mostly an increase in short-range contacts (**Figure 5D**). The overexpression of Topo III rescued the viability of the *parE^ts^* strain at non-permissive temperatures (Nurse et al., 2003, **Figure 5A**). The Hi-C contact map of a *parE^ts^* strain overexpressing Topo III at non-permissive temperature displayed a reduction in short-range contacts along the diagonal, but little changes regarding butterfly wings and mid-range contacts (**Figure 5E and F and Supplementary Figure 4C and 4D**). These observations demonstrate that short-range contacts are mediated by a topological structure that can be removed by Topo III, most likely precatenanes with single strand regions near the replication fork. Since butterfly wings, mid-range contact and terminus insulation are not suppressed by Topo III overexpression, they appear as a direct consequences of Topo IV inactivation and might therefore reflect the catenation of fully replicated molecules, a substrate impossible to unlink for Topo III (**Figure 5G**).

**Figure 5:**
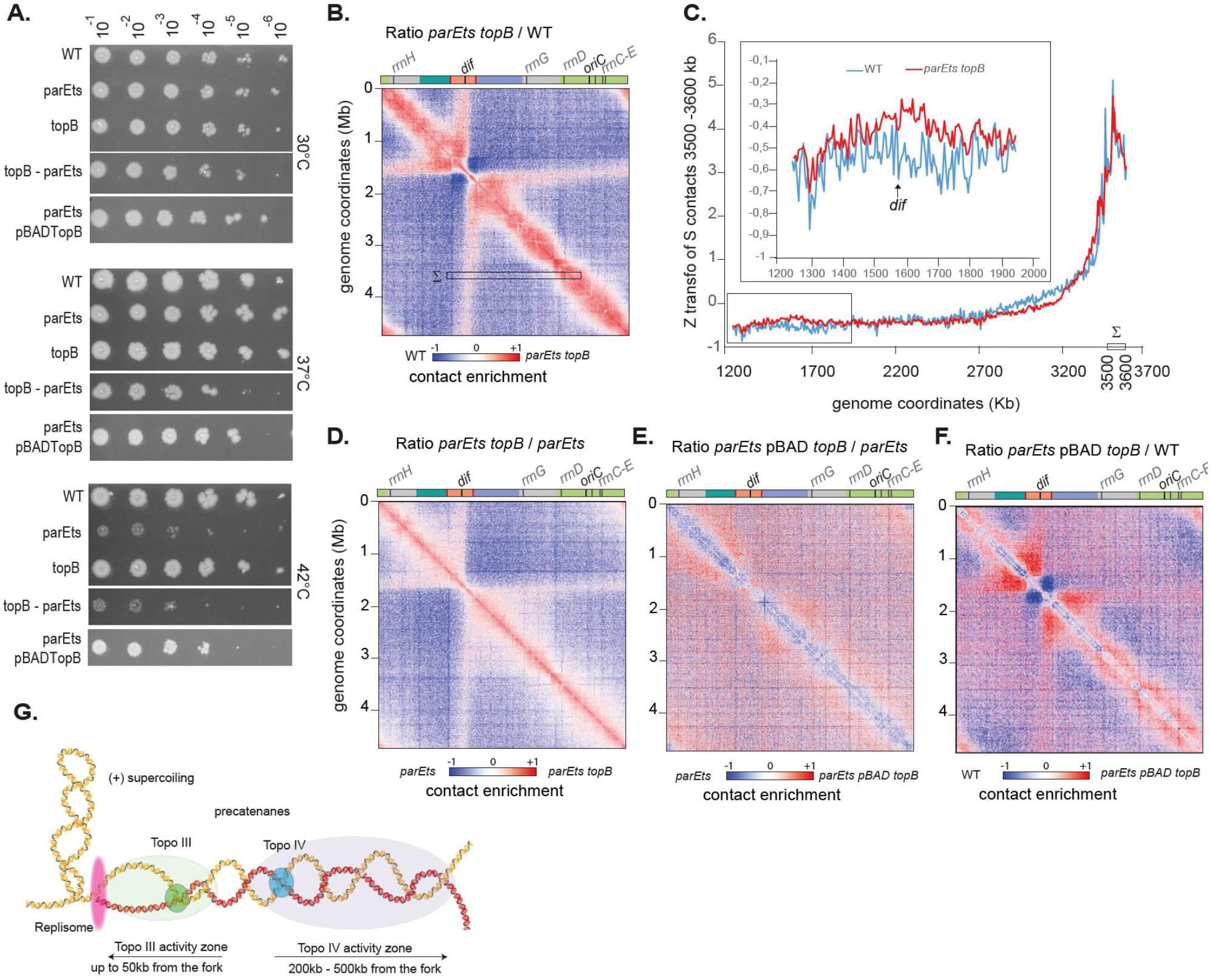
Topo III activity partially rescues Topo IV deficiencies on chromosome conformation. (A) Viability assay performed by droplet CFU of the WT, parE^ts^, parE^ts^ topB, topB, and parE^ts^ pBAD topB at 30°C, 37°C and 42°C. (B) Ratio of normalized contacts map binned at 5kb of for the parE^ts^ topB vs WT matrix. Macrodomains and interesting positions of the genome are indicated above the plot. The y axis indicates the genomic coordinates. A decrease or increase in contacts in the mutant cells compared with the control is represented with a blue or red colour, respectively. White indicates no differences between the two conditions. (C) Z transformation of the contacts made by a 100 Kb region located between the coordinates 3500 Kb and 3600 Kb (marked Σ in B) with the left replichore (coordinates 1250 - 3700 Kb). WT and parE^ts^ topB normalized matrix were analyzed. Inset, zoom on the terminal part of the genome. (D) Ratio of normalized contacts map binned at 5kb of for the parE^ts^ topB vs parE^ts^. (E) parE^ts^ pBAD topB vs WT. (F) parE^ts^ pBAD topB vs parE^ts^. (G) Graphic representation of the decatenation activity zone of Topo IV (blue) and Topo III (green) after the replication fork (pink). Sister chromatids are represented by double helices in red (sister A) and yellow (sister B). Topo III is able to act directly behind the replication fork at a 50kb resolution. Precatenanes beyond the Topo III decatenation activity zone, are dealt with by Topo IV which can act between 200kb and 500kb away from the replication fork.

### The position of butterfly wings is determined by MatP/*matS*, the Ter macrodomain organizer

The peculiar positioning of the butterfly wings at the edges of the Ter macrodomain **(Figure 3A)** prompted us to assess the influence of MatP on these structures. In the absence of MatP and at *parE^ts^* non-permissive temperature, the butterfly wings were replaced by a large region displaying increased long-range contacts (up to Mb distances), covering the entire *terminus* macrodomain **(Figure 6A).** The ratio plot of *matP parE^ts^* and *matP* contact maps further show an enrichment in mid-range contacts within the *terminus* macrodomain in absence of MatP (**Supplementary Figure 5A-D)**. This observation confirms that the features revealed by Topo IV inactivation are structurally connected, with MatP being responsible for the positioning of the discrete boundaries limiting the entry of putative precatenation links into the *terminus* macrodomain. These borders define the characteristic butterfly structure and favors the emergence of the terminus pattern.

**Figure 6:**
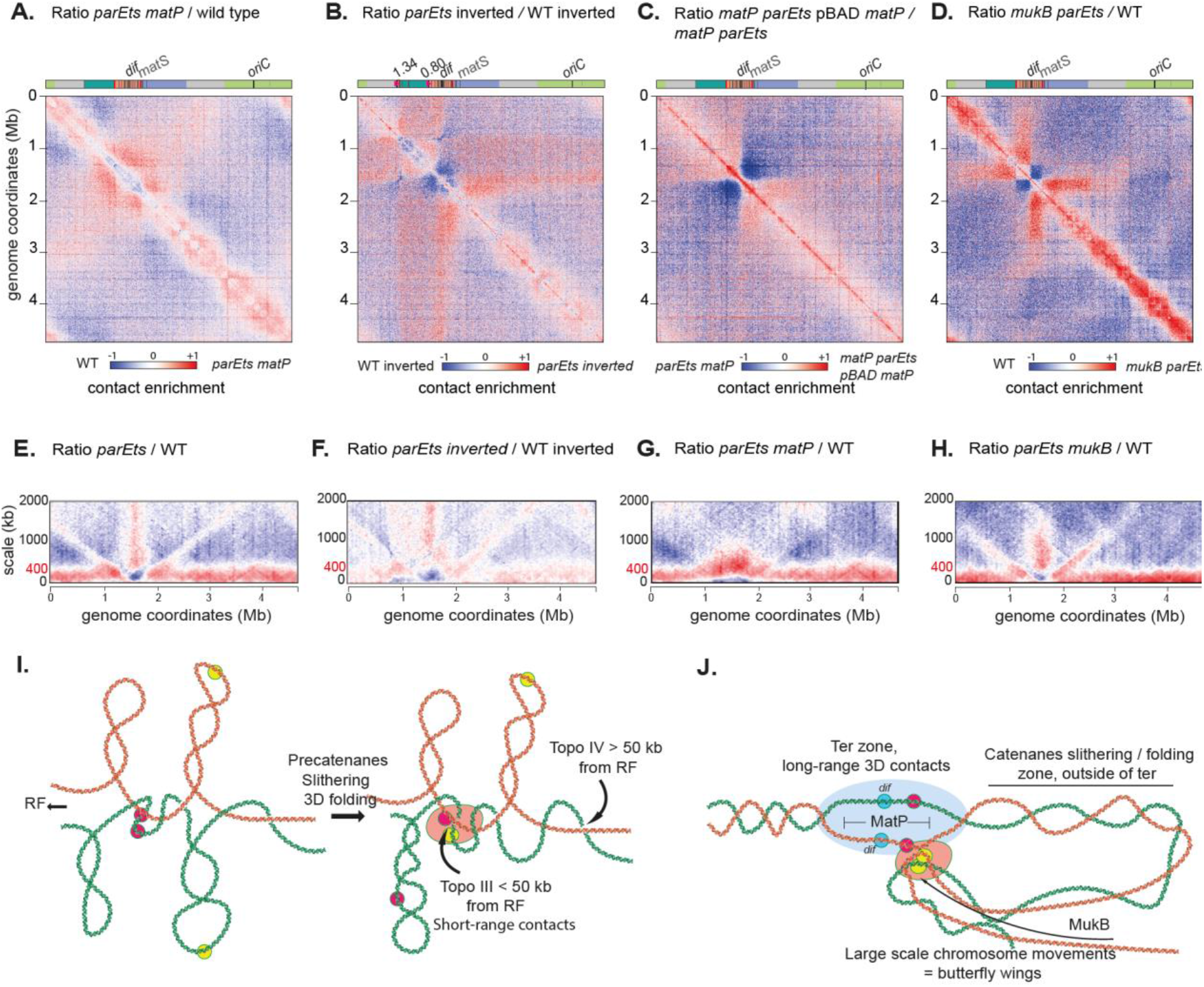
The butterfly position is determined by MatP/matS, the terminus macrodomain organizer and the bacterial SMC MukB. Ratio of normalized contacts map binned at 5kb, of (A) parE^ts^ matP vs WT, (B) parE^ts^ inverted vs WT inverted. The inversion was performed via the attR-attL system and inverted the region between 0.8Mb and 1.30Mb resulting in the displacement of matS sites. (C) parE^ts^ matP pBAD matP vs parE^ts^ matP, (D) parE^ts^ mukB vs WT. Scalogram ratio of normalized contact map comparing two conditions for each 10-kb bin along the chromosome: (E) parE^ts^ vs WT, (F) parE^ts^ inverted vs WT inverted and (G) parE^ts^ matP vs WT, (H) parE^ts^ mukB vs WT. Macrodomains and interesting positions of the genome are indicated above the plot. The y axis indicates the genomic coordinates. A decrease or increase in contacts in the mutant cells compared with the control is represented with a blue or red colour, respectively. White indicates no differences between the two conditions. (I) Graphic representation of putative sister chromatid precatenanes organization, dynamics and homeostasis based on Hi-C data of Topo IV and Topo III mutants. Two pairs of sister loci were represented in pink and yellow respectively. RF stands for replication forks. J) Graphic representation of the terminus region and the chromosome folding events that take place around it when decatenation is impaired. This region may function as a hub where unresolved entanglements of the two sisters formed during chromosome replication enter by a yet unknown mechanism to be separated by Topo IV. The low decatenation capacity in the parE^ts^ and parC^ts^ strains increase the persistence/recurrence of these events and their detection with Hi-C and imaging. MukB defines the maximum distance of loci able to contact the decatenation hub.

Based on these observations, we hypothesized that displacing the terminus macrodomain should reorganize the Topo IV-dependent structural features. We used bacteriophage λ site specific recombination to generate an inversion that moved the region associated with the base of the right butterfly wing (coordinate: 1. 342 702 Mb) to a region now 540 Kb upstream the right replichore (coordinate 0. 806 549Mb) (Esnault et al., 2007) **(Figure 6B, Supplementary Figure 5E and F).** This inversion resulted in the repositioning of five *matS* sites (matS 1-5) to a region upstream within the right replichore, which in turn resulted in the generation of a large chromosomal region (coordinates 0.8Mb to 1.3 Mb) devoid of *matS* sites and flanked by two *matS* regions. We observed a shift in the position of the right butterfly wing that coincided with the new position of the inverted *matS* sites. The butterfly wing was less precise, and long-range contacts appeared to extend from *matS*5 to *dif*. The positioning and strength of the second butterfly signal was not affected by the inversion, suggesting that the two butterfly wings are not functionally interlinked **(Figure 6B, Supplementary Figure 5E and F)**.

To test whether the butterfly structure is dynamic, we used an inducible *matP* gene and followed the formation of structures in the *terminus* region of the chromosome. After Topo IV inactivation in the absence of MatP, we induced the *matP* gene using arabinose. Thirty min of induction was sufficient to observe the reestablishment of the *terminus* pattern and preliminary butterfly wings **(Figure 6C, Supplementary Figure 5G and H).** These observations suggest that *parE^ts^* characteristic Hi-C signals correspond to dynamic structures, that can redistribute upon MatP binding to *mat*S.

### MukB controls the shape of the butterfly wings

The presence of MatP in the *terminus* macrodomain inhibits MukB activity in *WT* cells (Nolivos et al., 2016). In the absence of MatP, MukB is able to access the terminus region and has been shown to both accelerate the segregation of *terminus* loci (Nolivos et al., 2016) and change *terminus* conformation (Lioy et al., 2018). MukB also interacts with Topo IV and modulate its activity (Hayama and Marians, 2010; Li et al., 2010). We therefore tested whether MukB affects the patterns observed when Topo IV is inactive. A *mukB* deletion in itself did not create butterfly pattern (Lioy et al., 2018),suggesting that disrupting the ParC-MukB interaction does not alter Topo IV activity in the same way as *parE* or *parC* inactivation does. When combined with *parE^ts^*, the *mukB* mutation did not abolish the formation of butterfly wings **(Figure 6D).** However, their characteristics in the double mutant differ from those in the single *parE^ts^* mutant. Comparison of the *parE^ts^ mukB* mutant with the single *parE^ts^* mutant revealed that the basal portion of the butterfly wings were reinforced in the absence of *mukB* (i.e. higher frequency of contacts) and their length is reduced **(Supplementary Figure 5I)**. In addition, we observed an increase of contacts between all regions within the butterfly, so that they now resemble a self-interacting domain, rather than a stripe **(Figure 6D).** Scalograms, i.e. the aggregation of contacts over various scales from each bin (Lioy et al. 2018), clearly show the variations in wing shapes and sizes in the mutant strains (*parE^ts^, parE^ts^ inverted, parE^ts^ mukB* and *parE^ts^ matP*) **(Figure 6E-H).** In the *parE^ts^* strain, butterfly wings extend from the terminus to the origin of the chromosome. In the absence of MatP or in the inverted strain, they nearly disappear. In addition, MatP prevents the formation of mid-range contacts in the terminus region (**Figure 6F**). Finally, in the absence of MukB the butterfly wings abruptly stop ~1 Mb from *dif* on both replichores. The absence of MukB results in the exclusion of the *oriC* region from the butterfly structure. The increase in short range contacts detected along chromosome arms when Topo IV is inactivated (presumably corresponding to the accumulation of precatenanes) did not change in the absence of *matP* or *mukB*. This suggests that Topo IV partners do not significantly impact precatenanes dynamics. Therefore, MatP is involved in the positioning of the butterfly pattern, while MukB determines the wing length and contact density (**Figure 6H**). Moreover, MatP itself and/or the resulting MukB relocalization might prevents the progression of precatenanes into the terminus (**Figure 6F**).

## Discussion

### Alteration of each Topoisomerase induces specific chromosome conformations

Although DNA topology plays an important role in bacterial chromosome folding and compaction, the consequence of topoisomerase alteration on these processes are not yet fully understood. We took the advantage of Hi-C protocol improvements in bacteria to analyze chromosome conformation upon alteration of each of the four Topoisomerase of *E. coli*. Inhibition of Gyrase activity results in a strong reduction of short-range contacts over the entire genome. This agrees with previous report (Le et al., 2013) and confirms that supercoiling homeostasis is an important driver of CID organization in bacterial genomes. Surprisingly, because they have opposite catalytic activities, Topo I inhibition also reduces short-range contacts. Topo III and Topo IV inhibitions have the opposite effects, as they both increase short-range contacts: lightly and homogeneously for Topo III, but much more significantly and with a robust patterning for Topo IV. We investigated in details the nature and determinants of this patterning.

### New chromosome contacts emerging from Topo IV inhibition are intermolecular contacts

Topo IV inhibition induces an increase of chromosome contacts in the 50 – 200 kb range chromosome arms with the exception of a large terminus region that behaves differently. We observed that these new contacts are only established in replicating cells carrying circular chromosomes. Moreover, the activity of Topo III, which appears at very short range (0 – 50 kb), limits their amount. Altogether, these results demonstrate that short – mid range contacts detected when Topo IV is inactivated are produced by precatenanes **(Figure 6I)**. Their amplitude, 200-300 kb, can only be explained if catenated sister chromatids slides or fold in 3D on top of each other on a 200-300 kb window. By contrast, Topo III only modulated short scale contacts, which agrees with the idea that soon after replication sister loci are well aligned (no more than 50kb offset). In the absence of Topo IV some of these precatenane links persist away from the replication fork, out of reach of Topo III action even when overexpressed. Therefore, a few tens of second after their replication, catenane links cannot reach the Topo III activity zone, and might be entrapped by topological barriers (**Figure 6I).** The comparison of wild-type replicating and non-replicating cells (**Figure 4B**) confirms that during a regular cell cycle Topo IV and Topo III sporadically let few catenation links all over the genome that will contribute to sister chromatid cohesion (Joshi et al., 2013; Lesterlin et al., 2012). Future Hi-C and sisterC (Oomen et al., 2020) experiments performed on cells with synchronized cell cycle should give important insights on the nature of these inter-sister contacts and their molecular determinants.

### Very long-range contacts emerged as a butterfly pattern at the terminus

The butterfly pattern observed in the absence of Topo IV is an atypical Hi-C feature; Butterfly wings involve contact between the *dif* area (200 kb surrounding *dif*) and the rest of the chromosome. These contacts don’t follow genomic distance law and are not impacted by cis barriers (**Figure 3**). The density of contacts inside butterfly wings is dependent on the topological status of the cell (Topo III effect) and on MukB and their position determined by MatP/*matS* (**Figure 6**). We propose that the butterfly pattern is the signature of a failing decatenation hub (**Figure 6J**). *dif* is the major Topo IV activity site during regular cell cycle, this suggests that a large number of catenation links are removed at this locus (El Sayyed et al., 2016; Hojgaard et al., 1999). By contrast, the rest of the terminus region presents few Topo IV activity sites, but two prominent Topo IV binding sites at 1.2 and 2.8 Mb (El Sayyed et al., 2016). The positions of these binding sites corresponds to the border of the butterfly wing bases. Butterfly wings correspond to 3D contacts between the *dif* area and the rest of the chromosome. However, contacts between distal butterfly wings’ regions were not enhanced (**Figure 3**). Since butterfly wings were more robust when Topo III was inhibited and less robust when Topo III was overexpressed, we postulate that they correspond to contacts between catenation links scattered over the chromosome arms and the *dif* area (**Figure 6J**). Imaging confirmed the possibility of such contacts between the *ori* and *terminus* region in a small portion of the cells in a population, respectively 8% in *parE^ts^* and 2% in *wt* cells **(Figure 3)**. Several mechanisms might drive these contacts. One possibility is that protein marking catenation links is involved in protein-protein contacts between *dif* and other regions of the chromosome. A DNA binding protein marking positive supercoils, GapR, has been identified in *C. crescentus* (Guo et al., 2018). A yet unknown equivalent of GapR for catenation links (catenation crossing are topologically similar to positive supercoils) might exist in *E. coli*. Future experiments will be require to test if YejK which interacts with Topo IV and binds DNA (Lee and Marians, 2013) or another catenane binding factor, could play such a role. Another possible explanation is that regions of the chromosome presenting excess of catenation links are excluded from the bulk of the genome and are then available to interact with the decatenation hub in the terminus. Since the absence of MukB changes butterfly wing density, one hypothesis is that catenane links cannot be extruded as plectonemic loops. Successive unsuccessful contacts between a crippled decatenation hub in the terminus area and links dispersed on the genome might create the butterfly pattern **(Figure 6J**). The role of such contacts for chromosome segregation remains unknown, however the observation of duplicated *ori* foci close to the *terminus* foci and close to mid-cell in the *parC^ts^* strain **(Figure 3N)** suggest that this localization may promote decatenation perhaps by concentrating the few remaining active Topo IV in a hub like structure.

### Chromosome conformation capture proposes a new read out of topological constraints

Most of our knowledge on the control of topological homeostasis by topoisomerases comes from plasmid DNA topology analyses by 1D or 2D gel electrophoresis or electron or AFM microscopy (Cebrián et al., 2015). Recently, molecular genomics methods were implemented to map topoisomerase activities or binding on chromosome (McKie et al., 2020). Hi-C is a popular method to study chromosome conformation, and a derivative to track intersister contacts was recently designed in eukaryotes (Oomen et al., 2020). The use of *loxP* recombination to reveal proximity of sisters (Lesterlin et al., 2012) or their catenation (Mariezcurrena and Uhlmann, 2017) is also a powerful tool to decipher local chromosome topology, with a genome-wide *loxP* recombination based assay recently adapted to survey sister proximity in *V. cholera* (Espinosa et al., 2019). Our results suggest that Hi-C can also offer a read-out of topological constraints. Coupling all these approaches together, along with imaging, will refine our understanding of chromosome conformation dynamics during the bacteria cell cycle and particularly the contribution of inter-sister contacts.

## Methods

### Strains

The strains used for this study can be found in Supplementary table 1. All strains are derived from MG1655. All strains were grown in minimal media A (0.26M KH2PO4, 0. 06M K2HPO4, 0.01M tri sodium citrate, 2mM MgSO4, 0.04M (NH4)2 SO4) supplemented with 0.2% of casamino acids and 0.5% of glucose. All strains were grown at 30°C. The strain containing the thermosensitive allele of *gyrB* (*gyrB^ts^*) was then shifted at 42°C for 20min; the strain containing the thermosensitive allele of *parC* (*parC^ts^*) grown at 30°C which is already non-permissive temperature; and, the strain containing the thermosensitive allele of *parE* (*parE^ts^*) was shifted at 42°C for 60min at most. Time course of *parE^ts^* shift was performed by growing the cells at 30°C before performing a shift for 10, 20, 30, 40, 50 and 60 min. Strains containing the expression plasmid pBAD were cultivated with 0.2% arabinose to induce the expression of the gene under the control of the promoter pBAD. For the strain containing pBAD *topB*, the arabinose was present for the entire duration of the culture; for the strain containing pBAD *matP*, arabinose was added after 30min of shift to 42°C.

### Drugs and antibiotics

Inhibition of Topo I was done with a 5min treatment with 50ng/μl of Topetecan (Subramanian et al., 1995). DL-Serine Hydroxymate (SHX, Sigma CAS Number 55779-32-3), an inhibitor of seryl-tRNA synthetase which triggers the stringent response and prevents new rounds of replication, was used at a 10mg/ml working concentration for 90min. The efficiency of the drug is checked by FACS. Rifampicin was used for 10min at a 100ng/μl working concentration to inhibit transcription.

### Fluorescence activated cell sorting analysis

The number of nucleoid was monitored using BD fortessa cytometer with a 488 nm argon laser and 515–545 nm emission filter at a maximum of 5000 event per second. A minimum of 100 000 cells were analyzed per time point. Calibration was done with the samples of the stationary phase of the *WT* and *parE^ts^* at 30°C. For all samples, approximately 10^8^ cells were fixed in 70% EtOH, washed, marked with propidium iodide (2-20μg/ml), washed again, and resuspended in sterile 1× PBS pH 7.2 prior to analysis. FCSalyzer software (https://sourceforge.net/projects/fcsalyzer/) was used for data analysis.

### Hi-C libraries

Hi-C libraries were generated as recently described in (Cockram et al., 2021). 30mL of culture was grown in Minimal Medium A supplemented with casaminoacids and glucose until OD600nm ~ 0.2. Protein-DNA interactions were cross-linked by the addition of 37% formaldehyde (3% final concentration) for 30 min at room temperature with gentle agitation. Crosslinking was quenched with 2.5 M glycine (0.4 M final concentration) for 20 min at room temperature with gentle agitation. Fixed cells were then collected by centrifugation (4000 x g, 10 min 4°C), washed once in 1x PBS and snap frozen on dry ice and stored at −80°C until use. To proceed to the digestion of the cells, pellets were thawed on ice and resuspended in 1.2mL of 1xTE + complete protease inhibitor (EDTA-free, Roche). Cells were transferred to a VK05 tubes containing glass spreads, and then lysed with precellys (V750; 5×30s). Uncross linked proteins were then solubilized by incubating the lysed cells with 10% SDS (0.5% final) for 10min at room temperature. 1ml of lysed cells was then added to the digestion mix (3ml of dH2O, 0.5ml of 10x NEBuffer1, 0.5ml of 10% triton-X100, and 1000U HpaII) and incubated 3h at 37°C. Digested cells were pelleted by centrifugation and resuspended in 398μl of water before being added to the biotinylation mix (10X ligation buffer without ATP; dAGTtp 3.3mM; biotine-14-dCtp 0.4mM; 50U klenow (NEB5U/μl)) and incubated 45min at 37°C with agitation. Ligation mix (10X ligation buffer without ATP; BSA 10mg/ml; ATP 100mM; 250U thermoFisher T4 DNA ligase) was then added to the biotinylated DNA and the mix was incubated 3h at room temperature with gentle agitation. Protein-DNA complexes were then reverse crosslinked by adding the 20 μl of EDTA 0.5M, 80μl of SDS 10% and and 100 μl of proteinase K 20mg/ml and incubating at 65°C overnight. DNA extraction is made by phenol-chlorophorm and precipitation with ethanol. DNA is washed with 70% ethanol, resuspended in 130 μl of TE and then treated with 20mg/ml RNAse for 30min at 37°C. Samples were sonicated using a Covaris S220 instrument to obtain fragments between 300bp-500bp, then purified by AMPure XP beads. The Illumina process was performed according to manufacturer recommendation, with are 12 cycles of amplification. The size of the DNA fragment in the libraries are checked on TAE 1% agarose gel and subjected to paired-end sequencing on an Illumina sequencer (NextSeq500-550 – 75 cycles).

### Generation of contact maps

Generation of contact maps was done with *E.coli* analysis pipeline developed by Axel Cournac (https://github.com/axelcournac/EColi_analysis). Briefly, reads recovered from the sequencing were aligned with bowtie2 on the reference *E. coli* genome (NC_000913), the two ends were merged and the reads were filtered to have a mapping quality strictly > 30. Then each read was assigned to a HpaII restriction fragment (fragment attribution), more than 80% of the reads on average are conserved as informative reads. The contact matrices files are then generated, binned at 2, 5 or 10kb and normalized through the sequential component normalization procedure (SCN (Cournac et al., 2012)).

### Ratio and Ratio plots

In order to visualize the differences between two contact maps, their ratio is calculated as described in (Lioy et al., 2018). Briefly, contacts made in the mutant map are divided by the contacts made in the control map for each loci. Increase of decrease of contact in the mutant compared to the control will appear red or blue respectively and no changes will appear white. To visualize the contact signal at a smaller scale we use the ratio plot representation, as described by (Lioy et al., 2018). Briefly, the ratio-plots summarize the differences of contacts between two conditions made by each bin with its neighboring bins, regardless of their orientation. For each condition, the average contacts made by a bin along the genome with bins upstream and downstream at increasing distances (from 5kb to 1000kb or to 2000kb, in 5kb increments), was computed The log2(ACmutant/ACwt) was then displayed as a heatmap.

### Viability assay

For each strains, cells were serially diluted in LB (10-1 to 10-8) and 1μl of each dilution is deposed on LB agar plates. Plates were incubated overnight at 30°C.

### DNA density analysis

Cells were grown using the same conditions as Hi-C. 1ml of culture of a control sample at 30°C and a sample after a 60 min shift at 42°C for each strain were fixed every 20min (t0-t60), using an equal volume of 1X PFA (37% paraformaldehyde + glutaraldehyde + 1X PBS). DAPI (0.5μg/ml) is added and the mixture is incubated 10min in the dark. Cells were then washed and deposited on microscopy slides containing a freshly made agarose pas (1X PBS + 1% agarose). Samples were observed with a Spinning disk (YokogaWa) W1 system on a Zeiss inverted confocal microscope with a 63x, phase objective. To avoid bleaching, each field of view was only observed in the phase contrast channel before acquisition. Using ImageJ, the density of DAPI fluorescence in each nucleoid was calculated as a proxy for nucleoid density.

### Loci positioning analysis

Cells were grown akin to Hi-C samples (above). 1ml of culture of a control at 30°C and a sample after shift at 42°C for each strain, were fixed every 20min (t0-t60), using an equal volume of fixation medium (3.7% formaldehyde, 0,006 %glutaraldehyde, 1X PBS). Cells were pelleted and resuspended in 100μl of fresh Minimal medium A. Three drops of 2μl are deposited on glass slides containing a freshly made agarose pad (1X PBS + 1% agarose). Samples were observed with a Spinning disk (Yokogawa) W1 system mounted on a Zeiss inverted confocal microscope at 630-fold magnification. Using MicrobeJ (https://www.microbej.com/, (Ducret et al., 2016)), the position of *aidB* and *fear loci* is the cell in each condition at t0, t20, t40 and t60 after shift are calculated. An average of 600 cells were analyzed. Heat maps of the foci position are automatically generated by MicrobeJ. Two color localization was performed with *gusC parS pMT1* (ter) and *aidB parS P1* locus (ori) tags. Cells were grown at 25°C in Minimal medium A supplemented with casaminoacids and glucose until culture reached OD = 0.1 and then cells were shifted to 30°C for 1h. Cells were observed live on agarose pad on a thermo-controlled stage with a Zeiss inverted epifluorescence microscope equipped with led illumination. Localization of foci was recorded with the ObjectJ plugin of ImageJ https://sils.fnwi.uva.nl/bcb/objectj/. An average of 600 cells were analyzed.

## Acknowledgements

We thank other members of the Espeli and Koszul groups, Sylvie Rimsky, Adrien Camus, Martial Marbouty, Théo Foutel Rodier for insightful discussions. Axel Cournac, Etienne Jean, Lyam Baudry, Vittore Scolari, Cyril Matthey Dorey, Remi Montagne for Computational analysis tips. We thank Ivan Matic, Arnaud Gutierrez and Wei Lin Su and Agnès Thierry for technical advices. We thank the CIRB imaging facility. We thank Jean-Yves Bouet, Pat Higgins, Frédéric Boccard and Ken Marians for strains and plasmids. This research was supported by funding from the European Research Council under the 7th Framework Program (ERC Grant Agreement 771813 to R.K.) and ANR (HiResBac ANR-15-CE11-0023-03 to R.K. and O.E).

## Author contributions

BC, CC, RK and OE designed research. BC, IBC and CC performed the experiments, BC, IBC, CC and OE analyzed the data. All authors interpreted the data. BC, CC, RK and OE wrote the manuscript.

## Data Availability

Sample description and raw sequences are accessible on SRA database through the following accession number: PRJNA730396

