## Supplementary figures and table for "Genomic contacts reveal the control of sister chromosome decatenation in *E. coli*"

##### Supplementary Figure 1

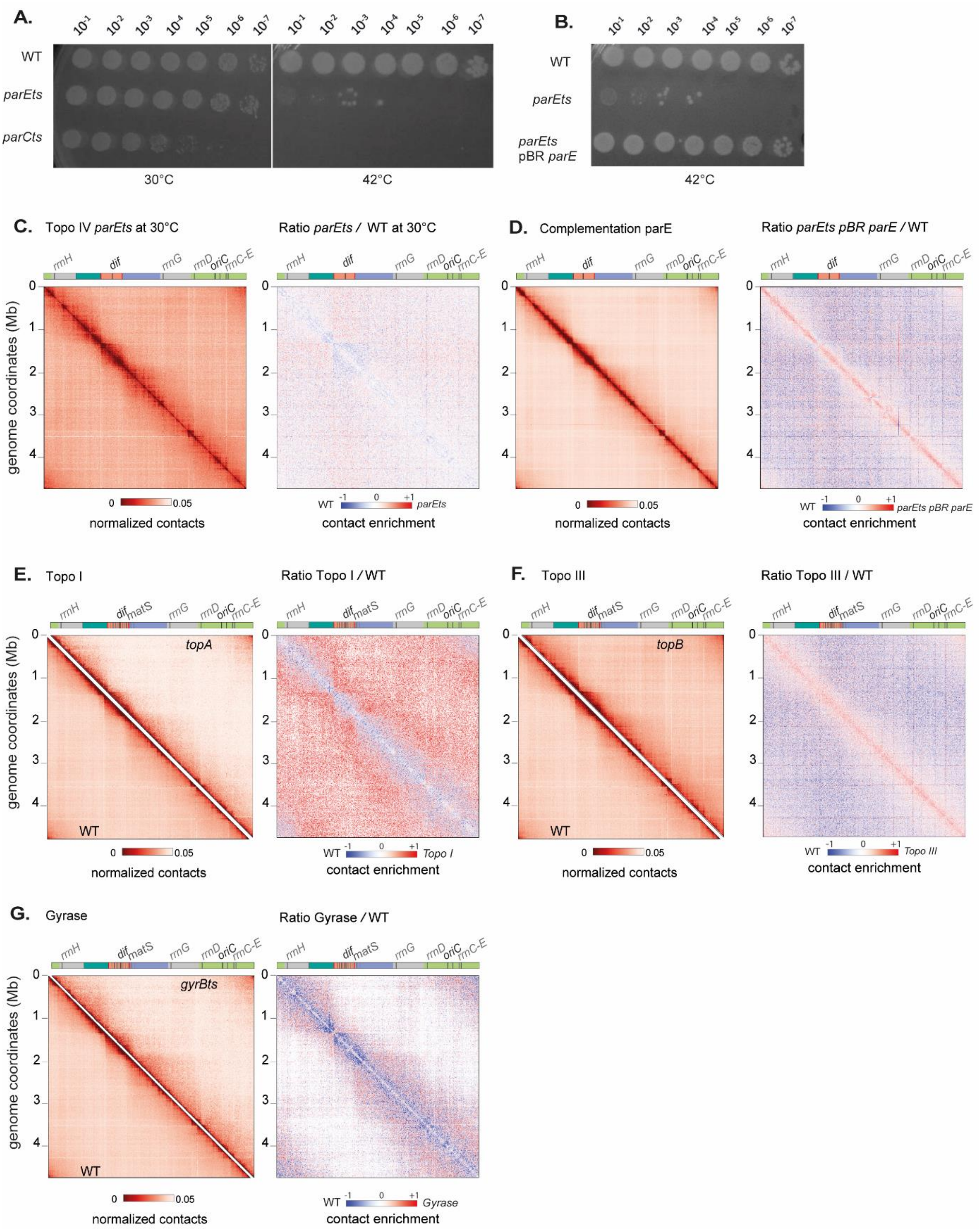

**Supplementary Figure 1**

(A) Viability in drop assay, of the WT, *parE*$^{ts}$, and *parC*$^{ts}$ at 30°C which is permissive temperature for *parE*$^{ts}$, while *parC*$^{ts}$ already displays defects and 42°C which is non permissive from both *parE*$^{ts}$ and *parC*$^{ts}$. (B) Viability in drop assay, of the WT, *parE*$^{ts}$, and *parE*$^{ts}$ pBR parE at 42°C. The expression of a functional ParE subunits restored the viability of the *parE*$^{ts}$ mutant at 42°C. (C) Normalized contact map of *parE*$^{ts}$ at permissive temperature (30°C) and the corresponding ratio of *parE*$^{ts}$ vs WT at 30°C. (D) Normalized contact map of *parE*$^{ts}$ complemented with pBR *parE* at non permissive temperature (42°C) and the corresponding ratio of *parE*$^{ts}$ vs WT at 42°C. (E-G) For each panels, symmetric halves of the normalized contact map binned at 5kb with the wild type (WT) on the bottom and the altered topoisomerase on the top, and the corresponding ratio matrix. (E) Topo I, inhibited by 5min treatement with Topotecan (F) *topB* deletion and, (G) *gyrBts* after 20min of shift to non-permissive temperature (42°C). Genome coordinates are indicated by the x and y axes. Interesting positions of the genome are indicated above the plot. *matS* sites are represented as gray bars. Macrodomain are represented by light green (ori), dark green (right), red (ter), blue (left), gray (NR/NL). For the normalized contact maps, the color scale of the frequency of contacts between two regions of the genome is indicated below (arbitrary units), from white (rare contacts) to dark red (frequent contacts). For the ratio matrices, a decrease or increase in contacts in the mutant cells compared with the control is represented with a blue or red color, respectively. White indicates no differences between the two conditions.

**Supplementary Figure 2**

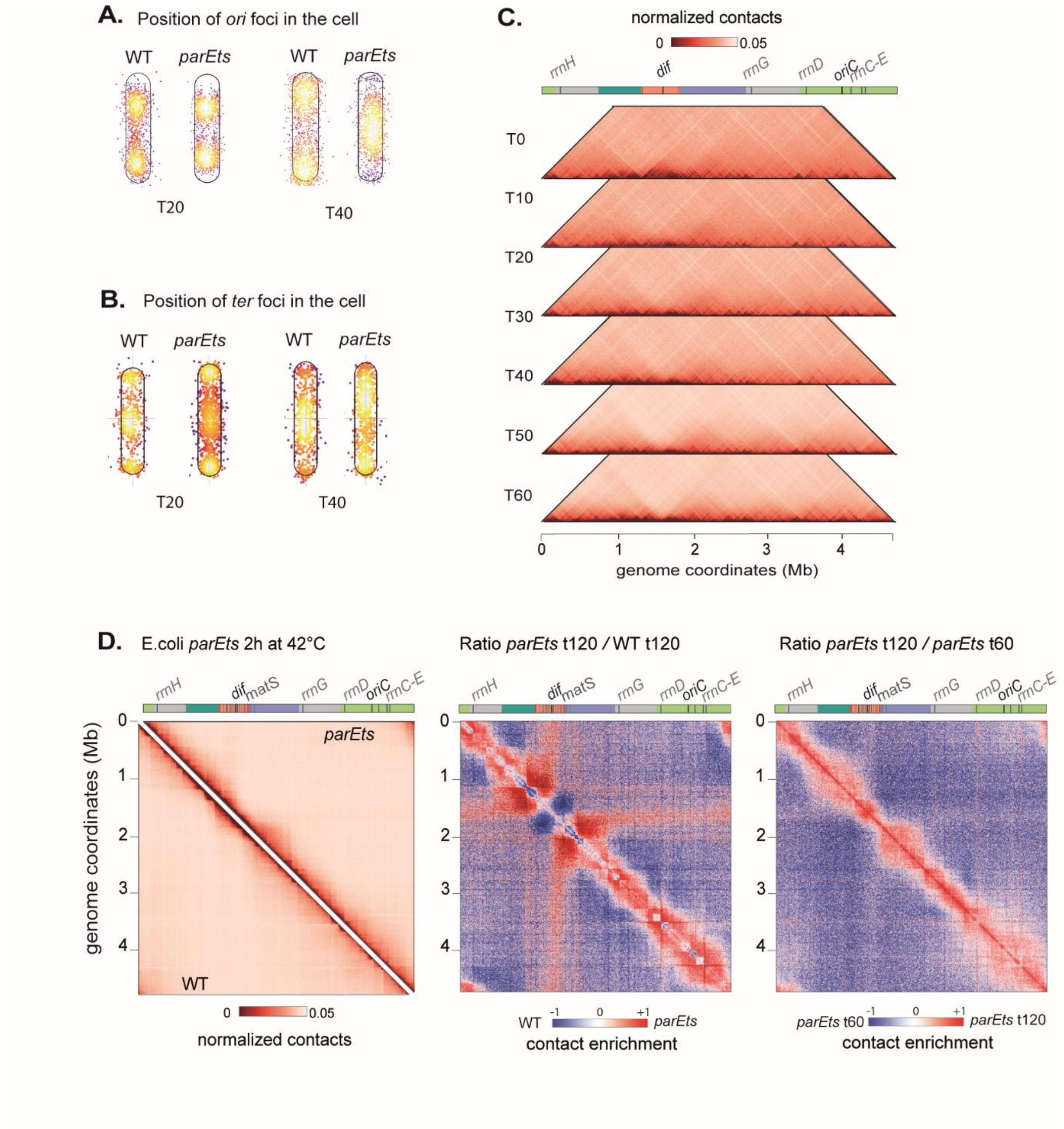

**Supplementary Figure 2**

(A-B) Average position of the *aidB parST1* (ori) foci (A) and *fear parST1* (ter) foci (B) in WT (left) and *parEts* (right) cells after 20min (T20) and 40min (T40) at non permissive temperature. Dots represents each detected focus (n=200) and the color scale represents the density of detected foci in the cell from black (0.05) to white (1). (C) Normalized matrices of the kinetics of the impact of Topo IV alteration with a time point every 10min after shift at non permissive temperature from t0 to t60. (D) On the left, symmetric halves of the normalized contact map binned at 5kb with the WT on the bottom and *parE*$^{ts}$ on the top 2h after shift at non permissive temperature (42°C) and in the center, the corresponding ratio. On the right, the normalized contact map comparing *parE*$^{ts}$ t120 to *parE*$^{ts}$ t60. Genome coordinates are indicated by the x and y axes. Interesting positions of the genome are indicated above the plot. *matS* sites are represented as gray bars. Macrodomain are represented by light green (ori), dark green (right), red (ter), blue (left), gray (NR/NL). For the normalized contact maps, the color scale of the frequency of contacts between two regions of the genome is indicated below (arbitrary units), from white (rare contacts) to dark red (frequent contacts). For the ratio matrices, a decrease or increase in contacts in the mutant cells compared with the control is represented with a blue or red color, respectively. White indicates no differences between the two conditions.

**Supplementary Figure 3**

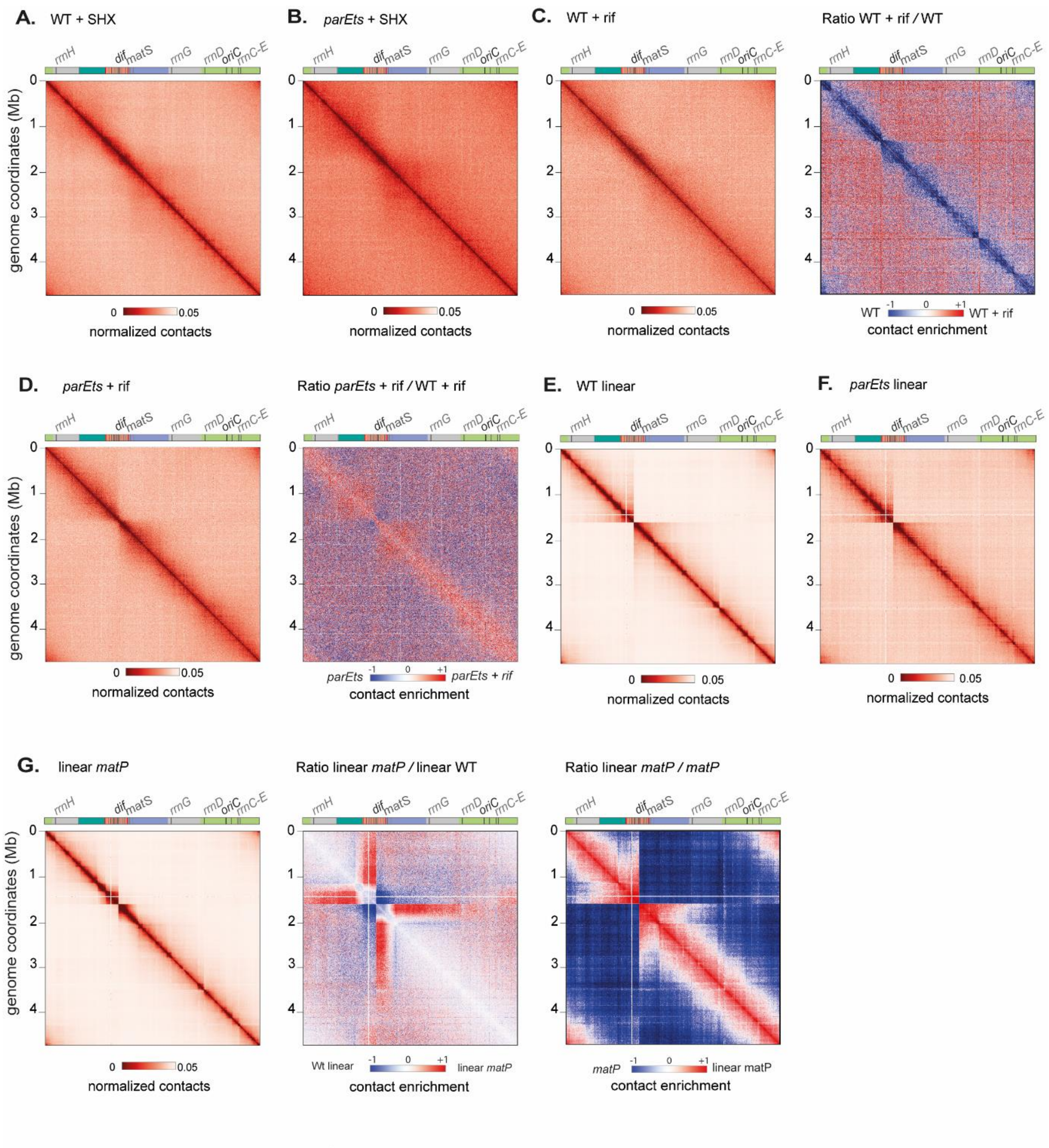

**Supplementary Figure 3**

(A) Normalized contact map binned at 5kb of WT treated with 10mg/mL of SHX for 90min at 42°C, (B) Normalized contact map binned at 5kb of *parE*$^{ts}$ treated with 10mg/mL of SHX for 90min at 42°C, (C) Normalized contact map binned at 10kb of WT treated with 100ng/µl of rifampicin for 10min, 50h after shift at 42°C, and the ratio of WT + rif compared to WT untreated, (D) Normalized contact map binned at 10kb of *parE*$^{ts}$ treated with 100ng/µl of rifampicin for 10min, 50h after shift at 42°C, and the ratio of WT + rif compared to WT untreated, (E) Normalized contact map binned at 5kb of *E. coli* with a linear chromosome 1h after shift at 42°C, (F) Normalized contact map binned at 5kb of *parE*$^{ts}$ in *E. coli* with a linear chromosome 1h after shift at 42°C. (G) On the left, normalized contact map binned at 5kb of *matP* in the linear *E. coli* strain. In the center, the ratio comparing linear *matP* to the linear WT and on the right, the normalized contact map comparing linear *matP* to *matP*. Genome coordinates are indicated by the x and y axes. Interesting positions of the genome are indicated above the plot. *matS* sites are represented as gray bars. Macrodomain are represented by light green (ori), dark green (right), red (ter), blue (left), gray (NR/NL). For the normalized contact maps, the color scale of the frequency of contacts between two regions of the genome is indicated below (arbitrary units), from white (rare contacts) to dark red (frequent contacts). For the ratio matrices, a decrease or increase in contacts in the mutant cells compared with the control is represented with a blue or red color, respectively. White indicates no differences between the two conditions.

### Supplementary Figure 4

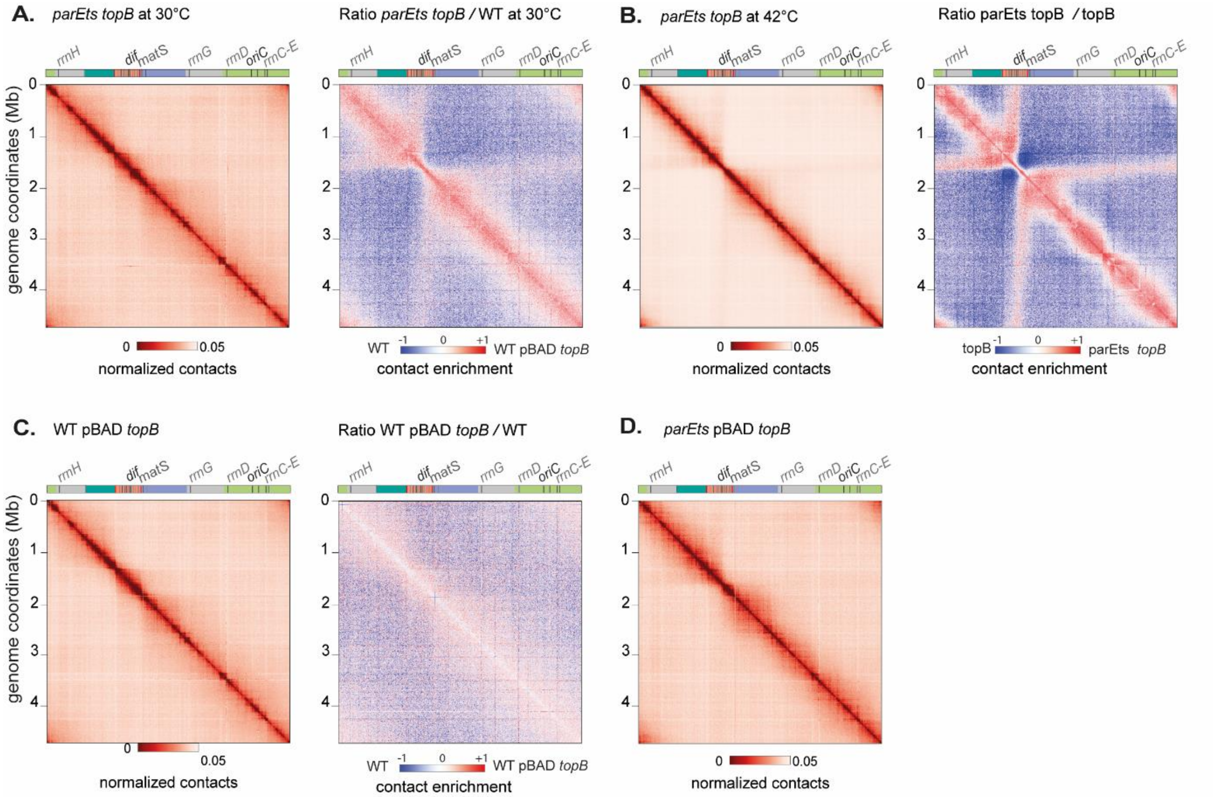

**Supplementary Figure 4**

(A) Normalized contact map binned at 5kb of *parEts topB* grown at 30°C and the ratio of the mutant compared to the WT. (B) Normalized contact map binned at 5kb of *parEts topB* 1h after shift at 42°C and the ratio of the *parE*$^{ts}$ *topB* compared to *topB* (normalized contact map of this mutant in Supp Fig 1I). (C) Normalized contact map binned at 5kb of pBAD *topB* grown at 30°C and the ratio of the pBAD *topB* compared to the WT. (D) Normalized contact map binned at 5kb of *parEts* pBAD *topB* grown at 30°C. Genome coordinates are indicated by the x and y axes. Interesting positions of the genome are indicated above the plot. *matS* sites are represented as gray bars. Macrodomain are represented by light green (ori), dark green (right), red (ter), blue (left), gray (NR/NL). For the normalized contact maps, the color scale of the frequency of contacts between two regions of the genome is indicated below (arbitrary units), from white (rare contacts) to dark red (frequent contacts). For the ratio matrices, a decrease or increase in contacts in the mutant cells compared with the control is represented with a blue or red color, respectively. White indicates no differences between the two conditions.

**Supplementary Figure 5**

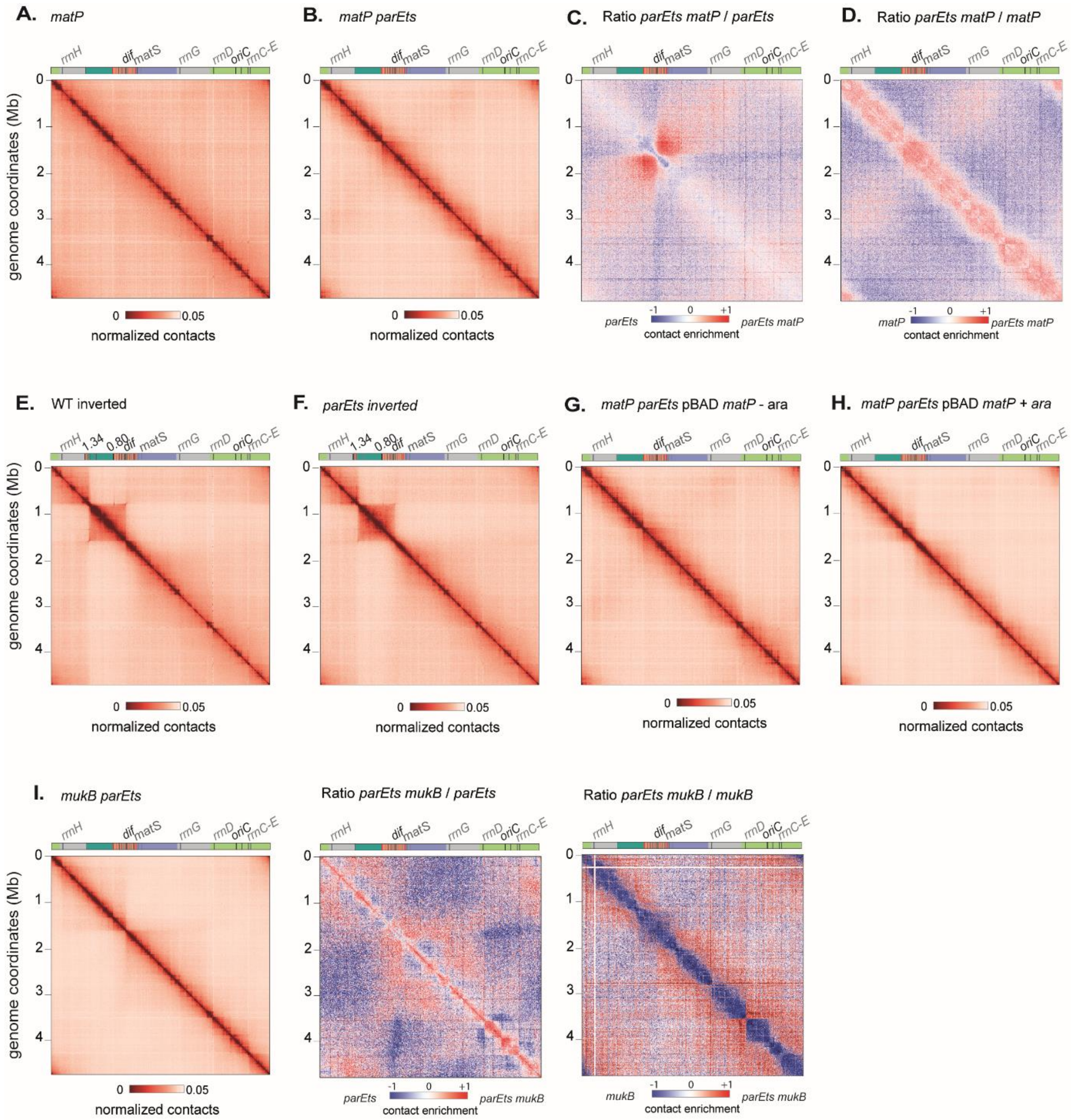

**Supplementary Figure 5**

(A) Normalized contact map binned at 5kb of *matP*, (B) Normalized contact map binned at 5kb of *matP parEts*, (C) Ratio of the normalized contact maps of *parE$^{ts}$ matP* and *parE$^{ts}$* 1h after shift at 42°C. (D) Ratio of the normalized contact maps of *parE$^{ts}$ matP* and *matP* 1h after shift at 42°C. (E) Normalized contact map binned at 5kb of WT inverted 1h after shift at 42°C, (F) Normalized contact map binned at 5kb of *parEts* inverted 1h after shift at 42°C, (G) Normalized contact map binned at 5kb of *parEts matP* pBAD *matP* non induced. Cells were grown at 30°C, then shifted for 60min at 42°C. (H) Normalized contact map binned at 5kb of *parE$^{ts}$ matP* pBAD *matP* induced with arabinose for 30min. Cells were first grown at 30°C, then shifted for 30min at 42°C before adding the arabinose for induction of the pBAD *matP*. (I) On the left, normalized contact map binned at 5kb of *mukB parEts* grown at 30°C and then shifted for 1h at 42°C. In the centre, ratio of the normalized contact maps of *parEts mukB* and *parEts* 1h after shift at 42°C. On the right, ratio of the normalized contact maps of *parEts mukB* and *mukB* 1h after shift at 42°C. Genome coordinates are indicated by the x and y axes. Interesting positions of the genome are indicated above the plot. *matS* sites are represented as gray bars. Macrodomain are represented by light green (ori), dark green (right), red (ter), blue (left), gray (NR/NL). For the normalized contact maps, the color scale of the frequency of contacts between two regions of the genome is indicated below (arbitrary units), from white (rare contacts) to dark red (frequent contacts). For the ratio matrices, a decrease or increase in contacts in the mutant cells compared with the control is represented with a blue or red color, respectively. White indicates no differences between the two conditions.

Supplementary Table 1: Strain used in this work

| GENOTYPE | REFERENCE OR SOURCE |
|---|---|
| **MG1655** | Espeli lab |
| **MG1655 *gyrBts*** | Espeli lab |
| **MG1655 *parEts*::Tc** | Espeli lab |
| **MG1655 *parCts*::Tc** | Espeli lab |
| **MG1655 topB::cm** | Nurse et al 2003 |
| **MG1655 *parEts*::Tc *ΔtopB*::Cm** | This work |
| **MG1655 *ΔyejK* FRT::cm ::FRT** | Lee et al 2013 |
| **MG1655 *ΔyejK* FRT::cm ::FRT *parEts*::Tc** | Espeli lab |
| **MG1655 *parEts*:TC + pBR-pBAD-*topB*:Kan** | Espeli lab |
| **MG1655 LR17 ptsACXI attlR attL (0.806549Mb/1.342702Mb)** | Gift from Frédéric Boccard |
| **MG1655 inverted *parEts* (0.806549Mb/1.342702Mb)** | This work |
| **MG1655 *ΔmatP*** | Espeli lab |
| **MG1655 *ΔmatP parEts*:Tc** | El Sayyed et al 2016 |
| **MG1655 *ΔmukB*** | Espeli lab |
| **MG1655 *ΔmukB parEts* ::Tc** | This work |
| **MG59** | Cui et al 2007 |
| **MG59 *ΔmatP*** | Espeli et al 2012 |
| **MG59 *parEts*** | This work |
| **MG59 *ΔmatP parEts*** | This work |
| **MG1655 *ΔmatP parEts* pBAD *matP*** | This work |
| **MG1655 *aidB* parS T1 pFH2973** | Espeli Lab |
| **MG1655 *parEts aidB* parS T1 pFH2973** | This work |
| **MG1655 *fear* parS T1 pFH2973** | Espeli Lab |
| **MG1655 *parEts fear* parS T1 pFH2973** | Espeli Lab |
| **MG1655 *aidB* parS P1 *gusC* parS T1 pFH2973** | Espeli Lab |
| **MG1655 *ygeB* parS P1 *gusC* parS T1 pFH2973** | Espeli Lab |
| **MG1655 *parCts aidB* parS P1 *gusC* parS T1 pFH2973** | This work |
| **MG1655 *parCts ygeB* parS P1 *gusC* parS T1 pFH2973** | This work |
| **Salmonella Typhimurium LT2** | Gift from Patrick Higgins lab |
| **Salmonella Typhimurium LT2 *parC281*(TS) zge-2393::Tn10** | Gift from Patrick Higgins lab |
| **Salmonella Typhimurium LT2 *parE206*(TS) zge-2393::Tn10** | Gift from Patrick Higgins lab |
